# Between group heritability and the status of hereditarianism as an evolutionary science

**DOI:** 10.1101/2023.12.18.572247

**Authors:** Charles C Roseman, Kevin A Bird

## Abstract

Hereditarianism is a school of thought that contends there are substantial evolved cognitive and behavioral differences among groups of humans which are both resistant to environmental intervention and are a root cause of differential social outcomes across groups. The relationship of betweengroup heritability 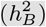 to within-group heritability 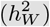 is one of the key theoretical components of hereditarianism and forms one of the bases for its claim to be an evolutionary science. Here, we examine the relationship between 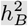and 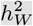 and its application to problems in the hereditarian literature from an evolutionary genetic perspective. We demonstrate that the formulation of the relationship between 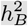 and 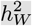used in the hereditarian literature has no evolutionary content. By re-writing the relationship between 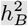 and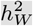 in a novel evolutionary framework, we demonstrate that there is no way to predict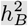 using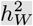 without considerable additional theory that is absent from the hereditarian literature. Furthermore, we demonstrate that the hereditarian technique that uses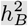 and 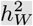 as a means of judging whether a given difference between groups may be plausibly ameliorated through environmental intervention is mathematically flawed. Lastly, we fill a gap in the hereditarian literature by writing out a means of using 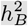 to predict the absolute difference between groups under a neutral evolutionary model and find that it is much smaller than claimed by hereditarians. In conclusion, we propose a path forward for the study of human variation that moves us past the ill-conceived nature vs. nurture question and allows us to focus on more productive issues.

## 1. Introduction

Following in the wake of the expansive post-genomic study of the genetics of complex traits, hereditarianism has caught a second wind. Hereditarianism is a school of thought in the psychological and social sciences positing that differences between groups of humans today in cognition, behavioral propensities, and economic outcomes are products of evolution and have a substantial genetic component that is largely insensitive to any environmental influence. Further, they propose these differences form the basis for sizable differences in life outcomes and the economic conditions among groups to a degree that makes human genetic variation a major causal factor in the course of human history (Jensen 1974; Rushton 1985; R. Lynn 1991; Rushton 1995; Rushton and Jensen 2005; R. Lynn 2008; Lasker et al. 2019; Lynn 2019; Fuerst, Shibaev, and Kirkegaard 2023).

As a field, hereditarianism draws heavily on the behavioral genetics tradition and, lately, genomics to address questions of group-level genetic differences in socially important traits in humans (Panofsky, Dasgupta, and Iturriaga 2021). Like their 20th century forbears, contemporary hereditarians make a point of emphasizing that theirs is an evolutionary approach (e.g., Jensen 1998; Winegard, Winegard, and Anomaly 2020; Kanazawa 2008). Where conventional evolutionary psychology hesitates to approach race- and other group-level differences, so their argument goes, hereditarianism embraces the full range of evolutionary theory to form a comprehensive science of human psychological and behavioral evolution (Jensen 1998; Winegard, Winegard, and A 2020; Rushton and Jensen 2005). Jensen (1998) characterized the hereditarian view as an evolutionary science by claiming

human individual differences and population differences in heritable behavioral capacities, as products of the evolutionary process in the distant past, are essentially composed of the same stuff, so to speak, controlled by differences in allele frequencies, and that differences in allele frequencies between populations exist for all heritable characteristics, physical or behavioral, in which we find individual differences within populations. (Jensen 1998, p. xii)

In the main, hereditarianism does not, however, use standard population genetic and evolutionary quantitative genetic theory to frame and investigate questions about the evolution of cognition and behavior. An important exception to this is in their use of the between group heritability 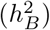 statistic to express the proportion of variation among groups that is attributable to evolved genetic differences (Jensen 1975, 1998; Warne 2020, 2021; Sesardic 2005; Fuerst, Kirkegaard, and Piffer 2021; Fuerst, Shibaev, and Kirkegaard 2023; Lasker et al. 2019). A key component of the Hereditarian 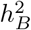 framework is the relationship between 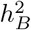 and within-group heritability 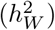, or the proportion of genetic variance within a group attributable to additive genetic effects (Lush 1947a, 1947b; DeFries 1972; Jensen 1998; Warne 2021). A core hereditarian position is that the magnitude of 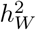, in combination with supplementary information, can be used to predict the extent to which environmental intervention will be able to eliminate cognitive and behavioral group differences (Jensen 1998; Sesardic 2005; Warne 2021; Winegard, Winegard, and Anomaly 2020; Fuerst, Kirkegaard, and Piffer 2021).

As Jensen argued:

Although one cannot formally generalize from within-group heritability to between-groups heritability, the evidence from studies of within-group heritability does, in fact, impose severe constraints on some of the most popular environmental theories of the existing racial and social class differences in educational performance. (Jensen 1975, p.1))

This use of the relationship between 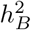 and 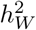 in this effort has been a point of frequent contention in the acrimonious disputes over the causes of disparate social outcomes of groups of humans for half a century (e.g., Jensen 1975; Feldman and Lewontin 1975). Conspicuously absent from the substantial literature on this issue during this lengthy span of time is any technical treatment of the theoretical relationship between 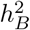 and 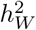 . Moreover, the issue of the relationship between 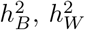, and the phenotypic response to environmental change at the group level is one of very few explicit engagements hereditarianism has with the core evolutionary sciences of population and quantitative genetics. As such, a critical examination of the relationship is a prime opportunity to evaluate whether the hereditarianism represents a genuine evolutionary perspective.

Here, we focus on the relationship of within-group variation to the differences among groups as a vehicle to critically examine the evolutionary and genetic claims of contemporary hereditarianism.

We show that the version of 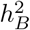1 used in hereditarian work is not evolutionary in any meaningful sense.^2^ We correct this shortfall in the hereditarian race science literature by writing out treatments of 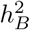 for use on evolutionary time scales under a neutral model. In doing so, we show that there is no relationship between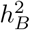 and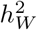 under an additive genetic model in which the effects of alleles are additive and insensitive to environmental context unless evolutionarily explicit models are used to relate them.

We further demonstrate that the hereditarian model that uses 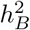 and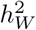 to assess the degree to which characteristics may be resistant to change in response to shifting environments has fundamental flaws. We then offer an empirical demonstration of the degree to which traits may change in response to environmental changes by drawing on results of studies of the quantitative genetics of, and historical change in, traits that have undergone secular changes in the recent past. These empirical results do not comport with hereditarian claims.

Moreover, while the hereditarian literature is focused on the *absolute* differences between groups in characteristics, it has not produced theory to relate that quantity to evolutionary genetic theory. We partly fill this gap by deriving an expectation for the absolute mean divergence between groups under neutrality thus providing a gauge of the magnitude of evolved differences among groups. Applying this framework to differences between groups of humans with a focus on populations of recent African and European origin or descent, we show that in the absence of natural selection we expect very small neutral evolved differences between groups for any trait even when assuming insensitivity of all genotypic effects to changes in environmental context.

We conclude by critically examining what heritability means outside of breeding contexts and outline a set of opportunities for behavioral and biological scientists to integrate the genetics of complex traits into the study of human behavior and evolution. We argue that placing the several traditions of the study of human variation on a genuinely evolutionary foundation has the potential to refocus the attention of scholars on more productive problems by dissolving and re-framing several of the controversies surrounding the causes of differences in behavior and social outcomes among groups.

## 2. Between group heritability and the hereditarian school of psychology

Jensen’s *How much can we boost IQ and scholastic achievement?* (1969) argued that behavioral and cognitive differences between socially defined racial groups in the United States have a substantial genetic (i.e., evolved) component. This claim attracted considerable interdisciplinary attention toward heritability and its implications for the causes of group differences in intelligence and the efficacy of educational and social policy. In an early attempt to place group differences in IQ into a population genetic framework, DeFries (1972), applied Lush’s (1947; 1947) formalism to establish a framework for studying the between-group differences in cognitive differences. DeFries concluded the among-race differences were likely to be minimally influenced by genetics. Jensen (1975) disagreed with DeFries’ rendition of the problem stating “[t]he [Lush-DeFries] formula is obviously only of theoretical interest, since we lack information on one of the parameters, *r*, the intraclass genetic correlation for the trait in question.” He would later revisit this same issue and reach a similar conclusion (Jensen 1998).

Feldman and Lewontin (1975) brought a formal population genetic perspective to the issue shortly thereafter (see also Jensen 1976; Plomin and DeFries 1976; Frankel 1976; Morton 1976; Feldman and Lewontin 1976). They noted that, while there is an algebraic relationship between 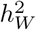 and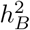 in that the terms on one side of the equation (the 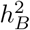 side) could be arrived at from those on the right side (the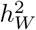 side). They claimed that it was obvious that there was no sense in which heritability at one level of analysis translated into heritability at another level in any genetically meaningful way in the Lush-DeFries formula. While correct, this point was not explained in much depth and has gone largely unheeded in the hereditarian literature.

The use of the Lush-DeFries equation in the hereditarian literature continues as the post-genomic era unfolds. In discussing their results from correlating estimated proportions of African and European ancestry on one hand, and IQ scores in a sample of self-identified African- and European-Americans on the other, Lasker et al. (2019) argue their results are consistent with 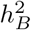 of IQ between 0.5-0.7 and supports Rushton and Jensen’s (2005) claim that 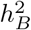 of IQ is between 0.5-0.8. The Lush-DeFries formalism was also recently employed to argue for a sizable genetic contribution to racial IQ differences (Fuerst, Kirkegaard, and Piffer 2021; Fuerst, Shibaev, and Kirkegaard 2023).

Warne (2021) revisited the issue of the relationship between 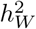 and 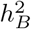 concluding that “The existence of the equation alone shows that the claim that ‘the genetic basis of the difference between two populations bears no logical or empirical relation to the heritability within populations’ (Lewontin 1970, p.7) is incorrect.” Based on this interpretation of heritability at different levels of organization, Warne echoed Jensen’s claim that there was a restricted range of possible environmental differences between groups defined as races in the context of the present day United States given plausible values for 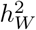 . He used this result to argue that some nontrivial proportion of the difference in IQ test scores between Black and white Americans must be genetic in some sense.

A core hereditarian claim is that the magnitude of 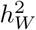 provides an indicator of the possible, or at least plausible, degree of environmental difference between groups. As Sesardic (2000, p.583), “[h]igh [within group heritability], together with some collateral empirical information, inductively establishes a non-zero [between group heritability]” . Sesardic condensed the hereditarian argument into a syllogism:

1. High 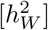 of IQ (among both whites and blacks).
2. Empirical data (mainly about the relation of certain environmental variables and IQ).
3. Non-zero 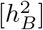.

(Sesardic 2000 p.587)

Notably, Sesardic’s version of Jensen’s hereditarian argument includes no theoretical basis for relating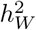 to 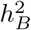 given some set of empirical observations.

Hereditarians are correct that there is *some* relationship between heritability within- and among-groups. After all, an evolutionary difference implies necessitates that within-group genetic variation be converted into between group variation. This relationship is an evolutionary one, however, and how the two quantities relate will depend a great deal on the particulars of the genotype-phenotype map and which processes of evolution are at work (Schraiber and Edge 2023). The hereditarian literature, however, does not provide a theoretical basis for evaluating which observations support a relationship between the two quantities.

## 3. Between group heritability in breeding contexts

Estimation of quantities representing the absolute and relative magnitudes of genetic and environmental variance in a population is a foundational part of modeling the evolution of complex traits and in crop and stock improvement (Falconer and Mackay 1996; Lynch and Walsh 1998). Likewise, human behavioral genetics has developed a parallel tradition to investigate the extent to which genetic and environmental effects might structure resemblances among relatives for psychological traits in a population (Downes and Turkheimer 2021). The formalism adopted by hereditarians (Jensen 1969, 1975, 1998; Lasker et al. 2019; Fuerst, Kirkegaard, and Piffer 2021; Warne 2021) was originally developed for animal breeding (Lush 1947a, 1947b) and adapted to the problem of among-group differences, (DeFries 1972), where “group” is usually construed as “race.” It is by no means clear, however, that applications of quantitative genetics meant to be used in breeding programs over the short term of a few generations translate well to problems over evolutionary time scales without substantial modification.

A challenge faced by breeders, evolutionists, and psychologists alike is the tendency for many organisms to associate in groups of varying degrees of relatedness and to share environments in ways that can confound genetic and environmental influences on phenotype (Falconer and Mackay 1996; Lynch and Walsh 1998; Walsh and Lynch 2018). For instance, family specific environmental effects may be highly correlated with genetic influences on offspring phenotype. Moreover, selection does not always operate at the individual level alone. It can operate at the among-family and among-population levels as well. Group and family selection are important for certain agronomic and animal breeding applications (e.g. Muir 1996).

Family selection represents an intermediate between individual and group level selection as conventionally understood (see Walsh and Lynch 2018, ch.21). Depending on the strength of environmental effects that are unique to families relative to the general effects of environment and genetic influences, a breeder might be better served by selecting on the basis of family membership alone, selecting desirable individuals from each family, or ignoring family membership entirely. Intermediate strategies using a combination of family and individual selection are also possible.

Lush (1947; 1947) developed a framework to evaluate which breeding strategy might be most effective when family-specific environmental effects may be strong thus making individual phenotype a poor indicator of breeding value. His model allows for a contrast to be made between the 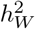 and 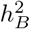, the relative values of which, properly scaled, may give a sense of when selecting within or on families will be more effective than conventional individual-level selection (Lush 1947b, 1947a; Walsh and Lynch 2018).

### 3.1 The Lush-DeFries equation

Lush’s aim was to build a general framework to decide whether a breeder would get better results by incorporating information about relatedness in their selection strategy (Lush 1947b, 1947a; DeFries 1972). He developed a technique for relating the differences at both the family and individual level to the response to selection. DeFries (1972) showed, under the assumption of infinitely large populations, we can relate the among-group heritability 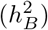 to within-group heritability 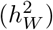 as

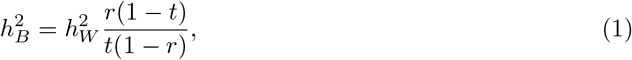

where *r* represents the intraclass genetic correlation within a group of individuals of a given degree of relatedness and *t* is the phenotypic intraclass correlation, which is a statistic summarizing the extent to which phenotypic values tend to cluster within groups as compared to the overall variation in the population. In this case, groups are usually construed to be sets of family members of a given degree of relatedness.

A difficulty with this rendition of the problem presented by the Lush-DeFries formalism is the fact that all of the quantities in this expression, save *r*, are functions of *r* and occasionally of one another. Lush (Lush 1947a, p.244) defines *r* as:

The numerical value of *r* is given closely enough for practical purposes by Wright’s coefficient of relationship if all the families belong to one breed, race or other population which was more or less freely interbreeding at a date only a few generations back, and if the relationship is computed to that date as a base.

The value of *r* is a function of the degree of relatedness separating two individuals such that, for example, full-sibs have *r* = 1*/*2, half-sibs *r* = 1*/*4, and first cousins *r* = 1*/*8 (Walsh and Lynch 2018, p.731). We can use *r* to predict the covariance among individuals of different degrees of relatedness arising from additive genetic effects as 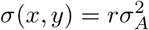 .

The phenotypic intraclass correlation (*t*) represents the tendency for phenotypes to cluster among siblings within a sibship, or any other set of group of individuals of like relatedness, and is a function of both genetic and environmental effects. In situations in which there are a large number of groups of relatives such as sibships with many siblings each, we can express the phenotypic intraclass correlation as

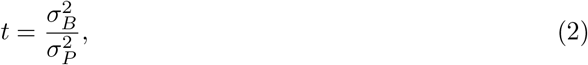

where 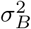 is the phenotypic variance among groups of individuals of like relatedness and 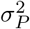 is the overall phenotypic variance (Lush 1947b, 1947a; Walsh and Lynch 2018).

The overall environmental variance 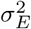 is the sum of the variance from effects common to members of a group 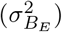 and that which reflects environmental effects that act on individuals irrespective of group membership 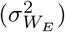:

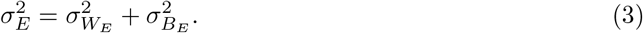

Note that while there is no general way to model the relationship between *r* and 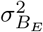, the environmental effects common to members of groups of a degree of relatedness is highly likely to differ substantially across classes of relatives, especially in mammals and other organisms requiring parental care and subject to parentally mediated environmental effects. We would not expect, for instance, for cousins (*r* = 0.125) to have the same environments as full siblings (*r* = 0.5) because full siblings are progeny of the same mother and, thus, will share common maternal effects. Other familial influences may have further effects. Given that we will doubtlessly encounter different magnitudes of 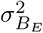 generic value of 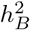 depending on which degree of relatedness we choose, we should not expect a across all degrees of relatedness in a population even at the same point in time.

We can decompose 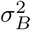 into 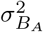 and 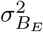, which are the among group and genetic and familial environmental variance — shared by members of a group of given relatedness to obtain the phenotypic intraclass correlation:

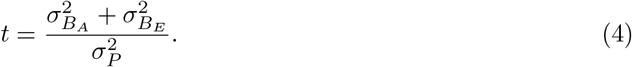

In this treatment, *t* is the proportion of phenotypic variance made up by genetic and environmental variance among groups of individuals of a given relatedness.

Unlike *r*, which is known for any type of relationship, *t* is dependent on the particulars of the causes of variation in the population. Moreover, *t* is in part a function of *r*. Since the degree to which relatives will resemble one another above and beyond what we expect by chance in the population depends, in part, on their degree of relatedness as represented by *r* and 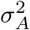, among other factors.

### 3.2 There is no evolution in the Lush-DeFries equation

Recall that Lush’s aim was to separate genetic and environmental effects within- and among-family allotments to understand the extent to which different breeding designs that took family structure into consideration, in one way or another, compared to those that were based on the standard practice of mass selection in which individuals were selected without regard for family membership (Lush 1947b, 1947a; Walsh and Lynch 2018). DeFries (1972) used the same general approach to build a framework to study differences among groups beyond groups of relatives as ordinarily construed in a breeding framework to problems in group differences on evolutionary time scales.

The first step in doing so is to define the within- and among- family components of additive genetic variance, which are

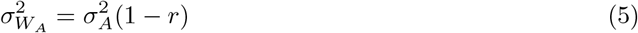

and

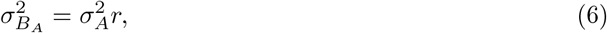

respectively. Since 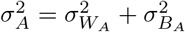, we can tell that 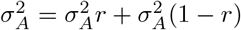.

There is nothing about the breeding process in and of itself that will effect a change in allele frequencies in a population (Hardy 1908) and changes in *r* do not represent evolution in the breeding context. They represent the investigators’ choice of a degree of relatedness in a random-mating population that is not evolving at any appreciable rate. As such, all we are doing is shifting our perspective on variation by changing our focus from one degree of relatedness in to another. In the absence of the action of evolutionary forces 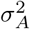 and allele frequencies will stay constant over a short period of time, irrespective of how we partition it, and we may partition it at will across sets of individuals of arbitrary relatedness given a large enough group.

As breeders, we would certainly have some control of the structure of relatedness in our population and would be able to choose groups of individuals of differing degrees of relatedness. Likewise, we might exercise some control over the values of 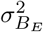 and 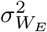 by way of cross-fostering offspring or even the implantation of embryos in unrelated dams (Atchley and Zhu 1997). In cases in which we had some sense of the prevailing norms of reaction, we might even be able to influence 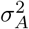 in our experiment, although this would be a haphazard exercise in all but the most well understood model organisms.

Ultimately, however, while 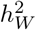 and 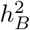 are both useful in deciding which selection scheme will be most efficacious to achieve an evolutionary end, they do not describe past evolution in the context in which they were originally proposed. As Lush put it:

For more complicated cases, such as selecting within a heterogeneous population containing different races, breeds or other groups whose ancestry has been separate for many generations, *r* is more difficult to measure. In such cases *r* is likely to have been augmented by the irregular incidence of mutations and by the accidents of sampling inherent in Mendelian inheritance. (Lush 1947a, p.244)

That is, the Lush-DeFries formalism as originally defined only applies to situations on short time scales and to those in which evolution can be ignored.

## 4. From the short-term to evolutionary time scales

The Lush-DeFries framework models only the disposition of variation at a given point in time. While this is useful for predicting how a population will respond to different combinations of family- and individual-level selection over short lengths of time, it relies on a period of minimal evolutionary activity to become useful for predictive applications. This issue can be resolved by recasting the Lush-DeFries equation in terms of population-level analogs of the genetic and phenotypic intraclass correlation.

### 4.1 Extending 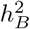 into evolutionary contexts

We can adapt the Lush-DeFries framework to evolutionary applications using a quantitative genetic analog of *F*_*ST*_ (called *Q*_*ST*_), which expresses the additive genetic variation apparent in a complex trait within- and among-groups as a proportion of all all genetic variance (Whitlock 1999, 2008; Whitlock and Guillaume 2009). It takes the form

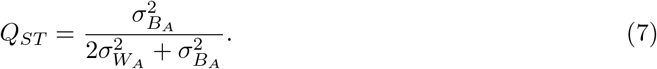

If a complex characteristic is evolving neutrally, we expect *Q*_*ST*_ for that trait to equal *F*_*ST*_ estimated from neutral genomic loci in the same organisms (Whitlock 1999). For the moment, we will focus on *Q*_*ST*_, which will allow us to be agnostic about which evolutionary dynamics led to the variation within and among groups. Our prossesually agnostic stance will allow us to evaluate whether there is a relationship between variation at the within- and between-group levels in the absence of an evolutionary model. Note that we assume that the environmental and genetic effects are uncorrelated throughout the remainder of the analysis.

Note that *Q*_*ST*_ closely resembles an intraclass correlation of the kind used in the Lush-DeFries equation. The doubling of 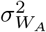 in the denominator is necessary to convert variance in breeding values/genotypes (i.e., zygotic variance) to variance in alleles (i.e., gametic variance). As demonstrated in Schraiber and Edge (2023) and implied in Edge and Rosenberg (2015, Eq. A1.4), the intraclass correlation in a neutral case may be written in terms of *Q*_*ST*_ or as

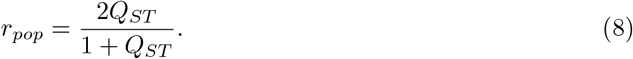

Substituting Eq. 7 into Eq. 8 and simplifying gives

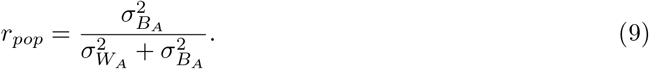

Replacing *r* in the Lush-DeFries equation with *r*_*pop*_ and writing all other out all other terms using their constituent variances yields

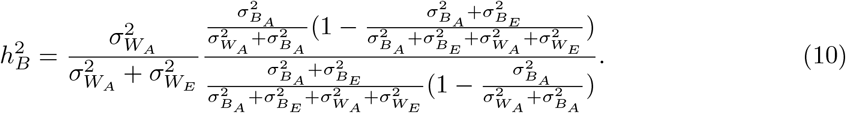

That Eq. 10 is a bit of a mess highlights one of the problems here. It features ratios of the products of ratios of correlated variables and developing an intuition about what happens to the system should any one of the quantities change is difficult in the extreme. The original Lush-DeFries equation sweeps most of the clutter under the rug, thus adding another layer of opacity to the problem.

The other problem with this rendition becomes obvious if we write 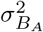 in terms of 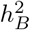as

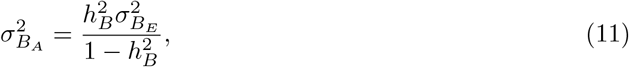

and expand 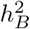 into its constituent terms to get

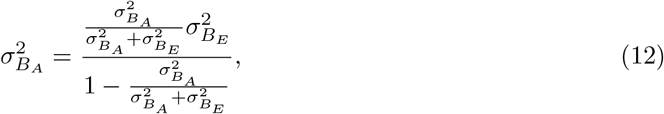

All of the within-group components cancel one another leaving only between-group components. There is also no unique solution to this equation for 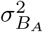 over the range of possible values that the terms might take. As such, 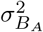 and 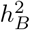 are not functions of any within-group components of variation and we require independent estimates of the quantities to do anything in the Lush-DeFries framework over evolutionary time scales.

As 1975 pointed out, there is nothing on the left side of the equation that is not already on the right side. The particular values of every constituent term of 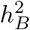 is either a function of or immediately derivable from information that we brought to the problem at the beginning. The Lush-DeFries equation is only informative in situations where the dog ate the first couple pages of your homework and you are left with only the intraclass correlations and 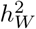 and need to produce 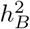 to salvage some partial credit. It follows from this exercise that there is no relationship between 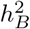 and 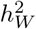 without considerable supplementary theory that is not present in the hereditarian literature. A similar conclusion was recently independently arrived at by Schraiber and Edge (2023).

### 4.2 The neutral case in terms of the coalescent

While re-casting the problem in terms of *Q*_*ST*_ provides a basis for performing the statistical operations over evolutionary time scales relevant to recent human diversification evolutionary version of the Lush-DeFries Equation, it does not provide a basis for a dynamical understanding of the evolution of group differences. The hereditarian literature tends to restrict the use of evolutionary theory to post-hoc applications of vague qualitative notions of the dynamics and outcomes of selection and adaptation. As such, we must look elsewhere for relevant theory.

In disciplines that form the core evolutionary theory, much of the genetics of the evolution of complex traits has been worked out for some problems while others remain the domain of ongoing efforts to build and empirically evaluate different models. In the latter category, work on theory about the evolution of complex traits is still working its way toward solutions that are good explanations of variation caused by a mixture of selection and other processes of evolution. In the former, however, the neutral theory of phenotypic evolution is well-developed and provides a useful basis for establishing how 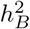 and 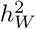 relate to one another in an explicitly evolutionary context.

Supposing neutrality, we can write *Q*_*ST*_ in terms of the amounts of time it takes alleles at a locus to share common ancestry within-or among- groups using coalescent theory (Slatkin 1991; Whitlock 1999). The average time it takes two copies randomly selected from across the entire species to share common ancestry (*τ*_*species*_) and the corresponding average time to coalescence within groups (*τ*_*within*_) give the amount of time that mutations have to occur such that they will produce differences within-or among-groups.

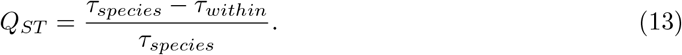

We can substitute this rendition into Eq. 8 above and simplify to obtain

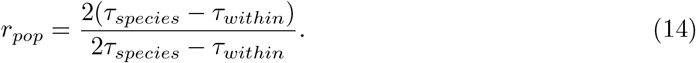

We may then write out the two varieties of heritability in terms of the coalescent times, environmental variance, and mutational variance as

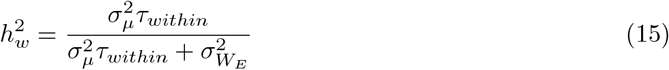

and

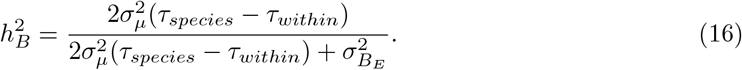

Recasting the Lush-DeFries equation in terms of coalescent times yields a predictably messy equation that we have relegated to Appendix 1. Suffice to say that the Lush-DeFries equation cast in terms of the coalescent does not yield unique soulutions for 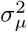, *τ*_*species*_, or *τ*_*within*_ except under circumstances in which 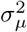 and *τ*_*species*_ = *τ*_*within*_. That is, it only works when there are no groups and no means of effecting evolution, which runs orthogonal to the original purpose of using the Lush-DeFries equation in the first place.

It shares a similar difficulty to the one encountered when examining the *Q*_*ST*_ rendition of 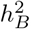 above. All of the constitutive mutational, drift, and environmental terms in 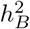 are already on the right hand side of Eq. 20 such that all within-group components cancel out leaving no dependency of 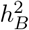 on the within-group specific terms. Thus, there is no relationship between 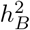 and 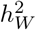 under a neutral model of evolution with additive and context-independent genetic effects (i.e., no epistasis or genotype by environment interaction).

This same point was made for the more general case of all evolutionary processes by Schraiber and Edge (2023). Moreover, by restricting our example to a neutrally evolving set of groups in which all genetic and environmental effects are uncorrelated we are presenting a best case scenario. Schraiber and Edge (2023) note that under certain models of evolution and in the presence of some kinds of genetic and environmental effects, when nonzero, 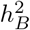 can take on any positive value, including those in excess of one.

Also noteworthy is how 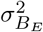 and 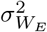 figure into this framework. While we can relate the strength of random genetic drift, a history of population-level relatedness, and the rate of the introduction of mutational variation into the population to the magnitudes of 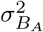 and 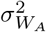, modeling changes in their environmental counterparts requires considerable supplementary theoretical work. Developments in cultural evolution (e.g., Uchiyama, Spicer, and Muthukrishna 2022) and the quantitative genetics of niche construction (e.g., Fogarty and Wade 2022).

### 4.3 To the degree that there is a relationship between *h*_2*W*_ and *h*_2*B*_, it is an evolutionary one

The principle lesson of all of this is the one Hardy expressed in one of the foundational documents of population genetics (Hardy 1908): There is nothing inherent to Mendelian rules of inheritance that causes evolution. As originally conceived, the Lush-DeFries equation assumes that no evolution is taking place. Supplementary evolutionary terms need to be included in a model to render it appropriate for the study of group differences on evolutionary time scales.

For instance, when modeling neutral divergence between independently evolving groups over short spans of evolutionary time we might adopt Lande’s 1976 expression relating 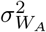 to 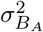. This takes the form 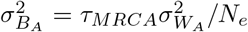 where *τ*_*MRCA*_ is time in generations since the groups shared a random mating common ancestral population and *N*_*e*_ is the effective population size, the inverse of which gives a sense of the strength of drift on a generation to generation basis. This could be substituted into Eq.10 to integrate it into an expanded Lush-DeFries framework. Other formulations based on the breeder’s equation or any of the many well-developed quantitative genetic models available to us would also apply.

Even with the addition of an evolutionary component to the between-group heritability setting, however, we would still need to address the environmental variance terms that are a part of the heritability statistics. As mentioned previously, there is little theory with which to structure investigations of this kind. Likewise, the relationship between 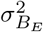 and 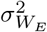 is not at all straightforward. This renders the second premise of Sesardic’s syllogism — “Empirical data (mainly about the relation of certain environmental variables and IQ).” (Sesardic 2005, p.587) — unable to sufficiently support his conclusion without considerable additional theoretical work to link these quantities.

## 5. Heritability and environmental differences between groups

The most prominent application of the relationship between 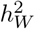 and 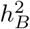 in hereditarian scholarship is in assessing the plausibility that a phenotypic difference between groups may be accounted for by an environmental difference (Jensen 1998; Warne 2020, 2021; Winegard, Winegard, and Anomaly 2020). The rationale for this approach is based on taking 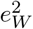, the environmental complement to 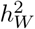, as an indicator of how strong an environmental influence on the between-group difference would have to be relative to the within-group environmental variation to cover the gap. So the hereditarian argument goes, if the magnitude of an environmental effect necessary to close the gap between groups is large relative to the within-group environmental variation, then it is less plausible, which is taken as evidence that there must be some nontrivial evolved component to the group difference.

While most of the applications of 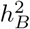 scores to issues in the hereditarian literature are post hoc and qualitative, Jensen 1998 proposed a framework to use 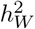 as a means to assess the plausibility of a range of 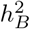 values even when direct estimates are not forthcoming. The strategy Jensen proposed is to scale differences between groups under the supposition of a given value of 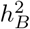 by the environmental proportion of within-group variation 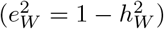.

Jensen used a simplified case in which within group variances for two groups are identical and scaled to unit variance and the phenotypic difference between the two groups is set to one within group standard deviation 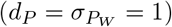(Jensen 1998; Sesardic 2000, 2005; Warne 2020, 2021). He expressed the environmental difference between groups as a function of 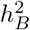 and 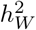 as

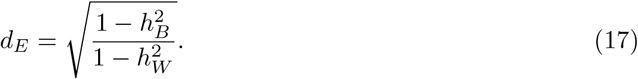

In Jensen’s own words, *d*_*E*_ is “[a] phenotypic difference between the means of two groups… expressed in units of the standard deviation of the average within-groups environmental standard deviation.” In this formulation, a *d*_*E*_ value between two groups with a high 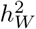 environmental effect would have to be larger to cover a given difference between groups when 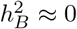 than in cases in which 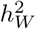 is low. Hereditarians take this to imply that 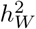 imparts, or at least hints at the presence of, a limit on the degree to which environmental effects might change group means.

Examination of Jensen’s framework for judging the plausibility of 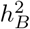 values given *d*_*P*_ and 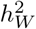 reveals some critical flaws that render it meaningless and without motivation.

### 5.1 Apples and oranges

Focusing on the denominator in Eq. 17, since 1^2^ = 1 and Jensen set 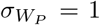, it is clear that 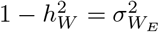 . This accords with the verbal description of *d*_*E*_ well enough as its square root gives the within-group environmental standard deviation.

A difference between the means of two groups (i.e., 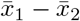), however, is not the between-group standard deviation. The between-group standard deviation is an expression of the mean distance of groups to the mean of the group means. Since the difference between the groups is set to 1, the absolute difference between either group and the mean of group means and the sum of square differences to the mean of means is 0.5. Thus, 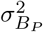 is 0.5 or 0.25, depending on whether we include Bessel’s correction (former value) or not (latter value) in the calculation of the variances in the two grou<u>p</u> case. The difference between means that would produce unit between-group variance is 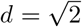. More succinctly, the Jensen equation conflates the between-group difference with the between-group standard deviation.

More importantly for our purpose of evaluating whether hereditarianism is based in an evolutionary theory, neither the genetic nor environmental difference between groups depend at all on the within-group components of variance in this framework except for the fact that the absolute among-group difference between groups is always held to be equal to the within-group standard deviation, which is standardized to unity. This fixes variation at arbitrary values and thus does not accommodate any dynamical account of change in variation. The only place where within-group variation enters the picture is as a standardization of the difference. This means that the constraint on between-group variation and is a separate arbitrary choice having nothing to do with any biological theory. As in the case in the relationship between *h*^2^ at different levels of organization, the hereditarian literature is devoid of theory that would explain why observed within-group environmental variation would be limiting of potential among-group environmental variation.

There are ways in which we might rescue something like Jensen’s equation. The simplest version would be to give 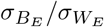, which would place the numerator and denominator on a commensurable scale. This does not, however, capture the spirit of Jensen’s verbal account of the quantity and does not relate 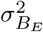 and 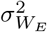 to one another in a theoretical framework.

Alternatively, we might couch Jensen’s equation in a directional selection framework. Suppose two groups, one that is at the ancestral mean and one that has been shifted to a new mean by some combination of selection and/or environmental shift. Further, say that all of the within group variational properties are identical. This would allow us to express the difference between groups as the product of a net selection differential and the within group heritability plus an environmental difference. This relates the within-group heritability to the between group difference through the breeder’s equation, but still leaves the relationship between observed 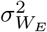 and potential 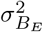 as an unexplored mystery.

### 5.2 Judging the plausibility of environmental differences using 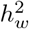

The Jensen equation is not suited for deciding if a difference between groups may be plausibly accounted for by environmental effects. Fundamental errors aside, it has no clear biological rationale because it is not based on any model of genetic effects in environmental context that might relate within- and between-group variational properties. Empirical investigation of secular change in trait values through time, however, may be useful in conditioning our expectations for the range of responses to environmental changes traits have experienced over short periods of time. Since evolutionary change may be ignored on the decadal time scale in most cases, all of the change through time in the mean of a characteristic will be environmental in origin. In this case, 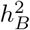 reflects cross-temporal genetic differences and is approximately nil on these generational time scales.

Table 1 gives estimates of the variational properties and secular changes on the decadal time scale in raw units and as standardized by *σ*_*E*_ for several traits that have undergone large secular trends over the past decades or centuries.^4^. We include the corresponding Jensen’s *d*_*E*_ values for each case for comparative purposes with the caution that they are meaningless.

**Table 1.** Caption

| Trait | $h^2$ | $\sigma_P^2$ | $\sigma_E^2$ | Secular<br>change<br>(/decade) | Secular<br>change<br>($\sigma_P$/decade) | Secular<br>change<br>($\sigma_E$/decade) | $d_E$ |
| --- | --- | --- | --- | --- | --- | --- | --- |
| Age at menarche (years) | 0.49 | 1 | 0.51 | 0.26 | 0.26 | 0.36 | 1.40 |
| Age at menopause (years) | 0.50 | 16 | 8.00 | 0.59 | 0.15 | 0.21 | 1.41 |
| Stature (cm) | 0.74 | 36 | 9.36 | 2.20 | 0.37 | 0.72 | 1.96 |
| Birth weight (g) | 0.31 | 237338 | 163763.22 | 36.80 | 0.08 | 0.09 | 1.20 |

While too small a sample with which to do much in the way of statistics, there are a few things we can draw from this exercise. For one, there is not a clear relationship between the *h*^2^ and and the secular change per decade scaled by *σ*_*E*_ or *σ*_*P*_ even though standardizing by *σ*_*E*_ imparts a positive correlation between the secular change and *h*^2^. This lack of a clear relationship is to be expected because there is no reason to believe that the magnitude of *h*^2^ will restrict or enable secular change via environmental influence.

Notable for our purposes is the fact that the fastest changing trait in terms of both the magnitudes of *σ*_*E*_ and *σ*_*P*_ is stature, which is also the most heritable by a considerable margin.^5^ Since the changes that motivate the secular trend are environmental in origin, these are realized changes and not changes of the hypothetical variety considered in the hereditarian literature. As such, these secular trends demonstrate that heritable characteristics can undergo much larger environmental changes than the hereditarian intuition would have us believe.

This illustrates the oft repeated, and oft ignored, caution that heritability and plasticity are not related to one another in a straightforward manner. Without a strong grasp of developmental and physiological mechanism, there is no way to predict how the means and higher moments of distributions of phenotypic characteristics in a population are going to change given a change in environments.

## 6. Expected neutral phenotypic differences among human groups

There is no necessary relationship between within- and between- group variational properties in the absence of an evolutionary model and the hereditarian literature offers no such model. Moreover, there are many evolutionary models from which to choose and which one will be most useful depends on the context and question. Where mainstream evolutionary theory tends to focus on changes in variances among groups and the magnitude of evolutionary distances, much of the hereditarian literature is concerned with the absolute differences between groups and it would be useful to have a simple model to condition our evolutionary expectations for this quantity.

To accomplish this, we adapt a neutral model of the evolution of complex traits to express the distribution of absolute pairwise differences between pairs of groups given that all evolution is effected by neutral mutation and random genetic drift. A neutral model has the dual appeal of being simple and providing a sense of what would happen if no natural selection was occurring.

The first step in building this expectation involves obtaining a value for 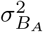 under the neutral expectation. Since under neutrality, *Q*_*ST*_ = *F*_*ST*_ and the distribution of evolutionary outcomes takes the form of a normal distribution, following (Whitlock and Guillaume 2009), we can rearrange Eq. 7 and substitute *F*_*ST*_ for *Q*_*ST*_ to obtain

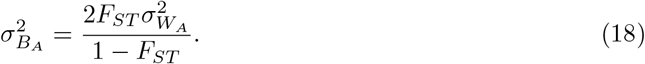

Estimates of neutral *F*_*ST*_ values can be found in population genomic investigations. Likewise, we can draw on the quantitative genetics literature for estimates of 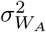 for whichever trait we wish to investigate. With these quantities in hand, we can write the expected absolute neutral additive difference between groups as

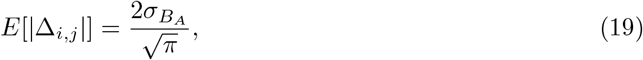

Details about this expectation and its corresponding cumulative probability and density functions may be found in Appendix 2.

### 6.1 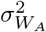 of cognitive traits, F_*ST*_, and neutral evolutionary differences between pairs of groups

To place differences between socially defined races of humans tentatively into a theoretical context on the time scale of the evolution of human genetic diversity, we can relate 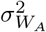 and *F*_*ST*_ to the absolute differences between groups. The human behavioral genetics literature tends to emphasize heritability over 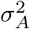 and does not often report, or attempt to garner, estimates of narrow sense heritability (*h*^2^) and instead focuses on one version or another of broad sense heritability (*H*^2^), which contains variance from a variety of non-additive genetic effects and interactions and covariance between genotypes and environments (Anholt and Mackay 2009). Some studies of individuals of varying degrees of relatedness and an increasing number of estimates derived from genome wide association studies are available, often in the population genetics and genetic epidemiology literatures. Likewise, with the advent of GWAS and their attendant polygenic indices, estimates of *h*^2^ have been obtained leveraging relationships between genotype and phenotype using loci from throughout the genome (e.g., Holland et al. 2020; Young et al. 2018).

Following Bird (2021) we use a range of narrow sense heritability values to bound the plausible span of 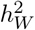 values obtained for educational attainment and cognitive test scores. These from 0.15 using a within family genomic regression technique that minimizes the upward biases inherent to the estimation of heritability in non-experimental contexts (Holland et al. 2020; Young et al. 2018), to 0.35 estimated using phenotypic resemblances among a wide range of classes of relatives (Chipuer, Rovine, and Plomin 1990; Cloninger, Rice, and Reich 1979; Devlin, Daniels, and Roeder 1997; Loehlin 1978; Rao et al. 1982), up to a doubtlessly overestimated 0.5 using twin studies and a genome regression technique (Polderman et al. 2015; Davies et al. 2011). These quantities are considerably lower than the usual reports of heritability in the human behavioral genetics literature because most exclude non-additive genetic effects and take measures to control for correlations between genotype and environment, epistasis, and other effects that tend to inflate estimates of heritability in twin studies and other sibling based family designs (Postma 2014). Multiplying these heritabilities by the customary variance intelligence test scores are forced to take 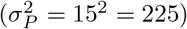 gives us our 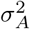 .

For our *F*_*ST*_ values between people of recent African and European ancestry, we adopt Bird’s (2021) value of *F*_*ST*_ = 0.12, which was estimated using loci from a frequency matched set of randomly chosen loci from the 1,000 Genomes Project (1000 Genomes Project Consortium et al. 2015). This value is typical of divergence between human groups from Africa and those in the remainder of the world (Alcala and Rosenberg 2017).

Figure 1 demonstrates that the bulk of the probability density of neutral absolute differences between pairs of groups is located at the very low end of the distribution. Unless there is pronounced natural selection acting to differentiate pairs of groups, large pairwise differences between groups would rarely occur.

**Figure 1.**
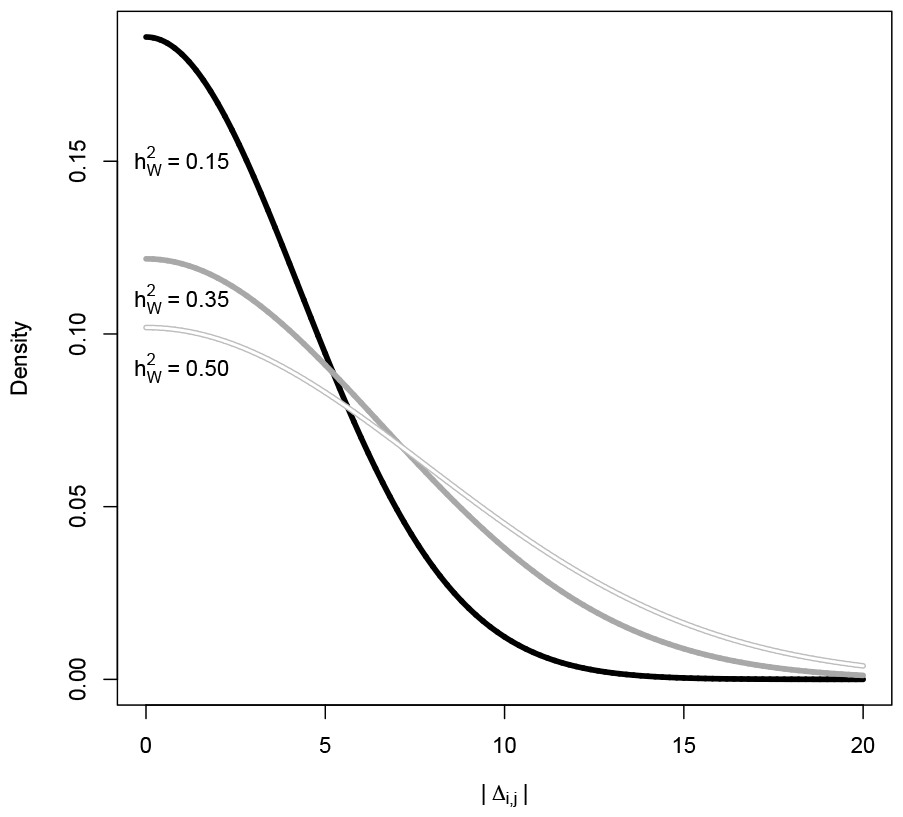
Probability density distributions for |∆_*i*,*j*_ | for 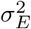 values of 0.15 (black line, 0.35 (grey line), and 0.50 (white on grey line). Note that the bulk of each distribution is located in regions of low divergence.

We wish to lay particular emphasis on the following point: *Under the neutral additive expectation, the difference can point either way because the differences are* ***absolute*** *and the expected difference between two lineages as sampled randomly after evolving under random genetic drift is 0*. As such, when we look at any of the differences between putative races reported in the hereditarian race science literature, we cannot expect an evolved difference between groups to resemble the phenotypic difference between groups without secure estimates of the magnitudes of among-group environmental effects, the degree to which there might be interactive effects (e.g., genotype by environment interactions and epistasis), and other quantities that are not presently in hand and are unlikely to be estimated anytime soon given the state of our understanding of canalization and phenotypic plasticity.

### 6.2 Related empirical & theoretical results

The only way that large amounts of evolutionary divergence among groups in IQ could be reconciled with the *F*_*ST*_ values estimated using neutrally evolving polymorphisms is if strong natural selection had acted to make the groups diverge from one another more than we might expect by neutral mutation and random genetic drift on their own. Depending on the number of and distribution of the effect sizes plus some other complicating factors (e.g. epistasis), natural selection would have caused small correlated shifts in the frequencies of many loci throughout the genome and other characteristic genomic signatures (Berg and Coop 2014).

The genomic evidence from several studies using distinct methods suggests that the loci underlying cognitive test scores do not show signs of having been under divergent natural selection between human populations. Using divergence metrics, Guo et al. (2018) was unable to reject the neutral hypothesis for a set of loci associated with educational attainment. Likewise, Bird (2021) found that measures of divergence for educational attainment and cognitive ability were indistinguishable from neutral background variation. Moreover, Racimo et al. (2018) used admixture graphing to attempt to detect natural selection on a set of loci associated with educational attainment and found a modest result in a sample from Asia, but none in samples from Africa or Europe. Even this limited positive result was unstable across different datasets Refoyo-Martínez et al. 2021. Recently, Howe et al. 2022 also did not detect positive natural selection on fluid intelligence or education in their British sample, finding non-significant correlations between SNP effect size estimates from their sibling GWAS and singleton density scores. Moreover, Zeng et al. 2018 did not find evidence of positive selection on intelligence or education in the UK Biobank using a Bayesian mixed linear model approach. If anything, the divergence among groups in the aggregate for loci underlying cognitive test scores and educational attainment are pedestrian when it comes to variation in the human genome or at even a lower magnitude than one would expect (Bird 2021). This is not likely to be simply a matter of lack of power to detect departures from neutrality as two of these studies found substantial deviations from the neutral expectation in other traits like height (e.g., Guo et al. 2018; Zeng et al. 2018; Howe et al. 2022).

Recent studies suggest that stabilizing selection might be the norm (Zeng et al. 2018, e.g.) thus keeping groups from evolving appreciably different cognitive abilities. While phenotypic differences among groups may not evolve much under stabilizing selection, the loci that contribute to their variation still evolve (Yair and Coop 2022). As Yair and Coop (2022) demonstrated, this continuing evolution of the loci influencing traits under selection can result in both a lack of portability of polygenic score indices across groups and an upward bias in the degree of phenotypic divergence among groups predicted using polygenic scores.

### 6.3 Challenges for hereditarians

At the outset of this section, we described this exercise of building a distribution describing the absolute evolved difference between groups as “tentatively” putting the problem on a theoretical foundation. We are probably getting ahead of ourselves in our framing of the problem. Given the lack of portability of polygeneic scores (Ding et al. 2023), growing evidence for the importance of genotype by environment correlation and interaction (Mostafavi et al. 2020; Cheesman et al. 2020; Durvasula and Price 2023), lack of evidence that cognitive ability scores are on an interval scale (e.g., Domingue 2014), and fundamental doubts about the legitimacy of legacy cognitive test score datasets (Sear 2022; Warne et al. 2019), there are much more basic scientific problems that need to be addressed before we can get too far down the road of rigorously comparing models and testing hypotheses about among-group differences in cognitive and behavioral traits.

The one actionable challenge hereditarians can take from this tentative exercise is that natural selection would have to have been quite strong to evolve differences on the order of scores of IQ points featured in the hereditarian literature (e.g., R. Lynn 1991; Rushton 1995; Rushton and Jensen 2005; Lasker et al. 2019). Efforts to improve the theory and method needed to detect departures from neutrality

## 7. Conclusion

We have demonstrated that the original Lush-DeFries framework is inapplicable to problems on evolutionary time scales, as Lush noted in 1947, because it assumes a single panmictic population in which the effects of mutation, selection, and drift are negligible. Expanding the Lush-DeFries framework to evolutionary problems by re-figuring the genetic intraclass correlation in terms of *Q*_*ST*_ demonstrates that there is no connection, much less dependence, between 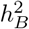 and 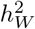 . Expansion of the Lush-DeFries equation into the its constituent variance clearly verifies the point made by Feldman and Lewontin (1975) that there is nothing on the right 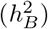 side of the equation that is not already on the left 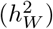 side. Moreover, the within-population variance terms drop out of the equation upon simplification demonstrating that 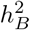 and 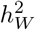 are distinct quantities. Our conclusions also agree with the more general result obtained by Schraiber and Edge (2023).

The only way in which 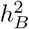 and 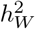 could depend on one another is if there was something about the operation of evolutionary dynamics that led to a dependency. For instance, a trait would have to be heritable within one or both of two groups at some point in the past for there to be evolution of group differences between them. While hereditarians hint at the need for such accessory theory to relate 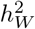 to 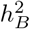, the hereditarian field has not produced any such theory. There are many different models that might apply in this situation, which means that hereditarians have all of their work ahead of them. To assist them in their efforts, we wrote out the relationship between 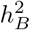 and 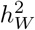 in a neutral evolutionary framework. In doing so, we demonstrated that there is no relationship between the two quantities under neutrality.

While we can relate 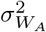 to 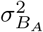 in a variety of ways using evolutionary quantitative genetic theory, the environmental counterparts of these quantities have no such theoretically informed interpretation. In the absence of considerable theoretical advances, the hereditarian assertion that, as Jensen put it, “the evidence from studies of within-group heritability does, in fact, impose severe constraints on some of the most popular environmental theories of the existing racial and social class difference” has no teeth.

Another core claim of hereditarianism is that 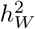 can be used as a means of judging the plausibility of hypotheses proposing that a difference between groups is entirely environmental (i.e., 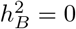. We show that Jensen’s equation used by hereditarians to evaluate these claims includes 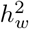 only as a scaling factor. There is no theory to model how 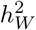 might impose any constraint on the possibility of environmental differences among groups. Moreover, we empirically demonstrated that high heritability is not an obstacle to environmentally induced phenotypic change. Highly heritable traits can exhibit secular change on the decade to century time scale.

Hereditarian claims about the degree of evolved difference among groups are not rooted in any evolutionary theory. We took a first step in placing the hereditarian hypothesis on a theoretical foundation by writing out the expected absolute difference between groups under neutrality assuming a model of heredity featuring additive genetic effects that are insensitive to environmental context. Under this simple model, we expect very small absolute differences among groups. Since loci associated with differences in cognitive traits and educational attainment almost always appear to be evolving neutrally even when those implicated in the variation of other traits do not (Guo et al. 2018; Bird 2021; Howe et al. 2022) and because there is every indication that a model of context-insensitive additive effects does not apply (Cheesman et al. 2020; Mostafavi et al. 2020; Hu et al. 2023), we suspect that even this neutral expectation overstates the possible difference. Whatever its size, any result of this kind is also irrelevant to understanding what a difference might look like in another social or environmental context.

### 7.1 Hereditarianism is not evolutionary

Much of the hereditarian literature is devoted to complaints about how hereditarian views are excluded from mainstream science because their questions and conclusions are held to be taboo for political reasons by mainstream scholars (Carl 2018; Winegard, Winegard, and Anomaly 2020). Setting aside the issue of whether hereditarianism is being marginalized for political reasons on account of its evolutionary perspective (cf. Jackson and Winston 2021; Bird, Jackson, and Winston, in press), we have demonstrated here that hereditarianism is not evolutionary. The models the field uses to approach questions about variation at the individual and group level have no explicit evolutionary content. This kind of study can be shoehorned into various evolutionary frameworks, as we have demonstrated here, but hereditarians have not taken the time to do so. This lack of connection between evolutionary theory and data through methods that are constructed in a theoretically informed way means the evolutionary statements in the hereditarian literature are products of nothing more than *post hoc* exercises in a-theoretical verbal interpretation.

This renders the hereditarian complaint of being excluded from mainstream social and evolutionary sciences on account of their application of evolutionary thought moot because there is no such application of evolutionary thought in the first place. This lack of evolutionary thinking also afflicts the hereditarian use of polygenic scores for comparisons among groups (see Bird 2021; Bird, Jackson, and Winston, in press, for a critical account.) and the use of polygenic scores and admixture analyses (see Kim et al. 2021; Schraiber and Edge 2023, for a demonstration of why this is).

### 7.2 What is heritability for?

When we turn the critical eye toward the mainstream study of human variation, several challenges and opportunities for the interdisciplinary study of human variation become apparent. Heritability carries with it almost no information about the genetic architecture, genotype-phenotype map, or propensity for the variational properties of a trait in a population to change in response to changes in environmental conditions. Given estimates of *h*^2^ and 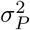, we gain very little information about the genetic architecture or genotype-phenotype map of the trait even if we can be certain that our estimates would allow us accurately to predict a response to directional selection should the environment remain much the same. We would not be able to tell how many genes were involved, beyond saying that there were probably a lot of them, nor could we offer informed comment on their patterns of interaction with one another and the range of environments to which individuals in the studied sample were exposed. That is, heritability estimates of any kind and their cultural evolutionary analogs are meant to describe the evolutionary dynamics of aggregates and have next to nothing to say about underlying mechanism without considerable additional work (Bookstein 2016). Aside from its status as a bycatch statistic genetically informed techniques used to disentangling the confounding of genetic and environmental effects in non-experimental settings (Downes and Turkheimer 2021) and its natural resonances with cultural evolutionary frameworks (Uchiyama, Spicer, and Muthukrishna 2022) it is not clear what heritability is doing for the social sciences.

This worry about the role of heritability is not restricted to the social sciences. The question “what is heritability for?” also troubles the mainstream study of the evolution of non-human organism. Developments in evolutionary genetics over the last few decades have shown that heritability is a poor indicator of how quickly a population will evolve by directional natural selection (Hansen, Pélabon, and Houle 2011). Moreover, heritability has several unpleasant statistical properties stemming from the fact that it is the ratio of correlated quantities and does not take into consideration issues of scale that are inherent to the evolution of complex traits (Hansen, Pélabon, and Houle 2011).

Evolvability, the mean square standardized additive genetic variance (Hansen, Pélabon, and Houle 2011), is a much more useful quantity when it comes to problems of how quickly a trait will evolve by natural selection. It gives the single generation response to selection relative to the trait mean should selection on the trait in question be as strong as that on fitness itself. While this is not an evolutionary speed limit, it serves as a useful benchmark for comparing the pace of evolution by natural selection across contexts.

A good example of this is endocranial volume (ECV) in humans. It tends to be highly heritable — however heritability is estimated — often exceeding 0.9 (Miller and Penke 2007). It is, however, less evolvable than other volumetric or mass-based characteristics in humans (Miller and Penke 2007). From a naïve heritability standpoint, it would be completely unsurprising that ECV might evolve quickly, but this would be illusory as the amount of additive genetic variation available is so restricted. An evolvability standpoint makes it clear that it would take very strong selection to effect a fast evolutionary response in ECV.

Since evolvability involves the estimation of the additive component of genetic variance, traditional twin designs will not be adequate to estimate evolvability in humans as the covariance among different classes of twins contains non-additive genetic components in addition to substantial environmental confounds (Falconer and Mackay 1996; Lynch and Walsh 1998). Moreover, psychological characteristics, with the exception of certain endophenotypes, are not expressed on ratio scales, which prohibits their meaningful expression through mean-standardization.

Since evolvability involves the estimation of the additive component of genetic variance, traditional twin designs will not be adequate to estimate evolvability in humans as the covariance among different classes of twins contains non-additive genetic components in addition to substantial environmental confounds (Falconer and Mackay 1996; Lynch and Walsh 1998). Moreover, psychological characteristics, with the exception of certain endophenotypes, are not expressed on ratio scales, which prohibits their meaningful expression through mean-standardization.

### 7.3 Moving forward

These considerations indicate that a promising avenue for future investigation might be a focus on resolving issues of trait characterization and measurement that plague phenotypic investigations whether they be behavioral, cognitive, physiological, or morphological. In this respect, the study of psychological characteristics is very much in the same boat as the study of any other class of characteristic. Evolutionary morphology, in particular, is producing considerable theoretical strides toward using the properties of scales of measurement to arrive at more meaningful evolutionary conclusions (Voje et al. 2023).

As suggested by Uchiyama et al. (2022), one site of potential synthesis that may prove to be exceptionally productive is at the nexus of evolutionary genetics and cultural evolutionary studies. Besides having anastomosing intellectual roots in population and quantitative genetics, these fields share compatible methodological frameworks.

As a cautionary note, it is also apparent that hereditarian investigations are not that different in conceptual structure and method from many other more mainstream scholarly traditions focusing on human evolution such as human biology and biological anthropology. Were we to take a hereditarian paper on race differences in intelligence test scores and replace “race” with “population” and “IQ” with “femur length” coupled with a few adjustments account for idiomatic differences between disciplines, it would be difficult to pick the redecorated hereditarian paper out from those in human biology and anthropology.

Much of the work in the traditional study of human variation tend to ignore population history and structure and are unconcerned with confounding of genetic and environmental effects (e.g., Stock 2023). We point this out not to give succor to hereditarians but to highlight the urgency with which the mainstream sciences of human variation should reform themselves. Hereditarians could make a positive contribution to this effort by focusing on basic evolutionary questions of the kind that all scientists of human variation face.

A process of reformation would provide a salutary change in the way in which research linking evolution to individual and group differences is conducted and argued. As we have demonstrated here, there is an enormous gap between the bedrock of evolutionary theory and the available data on one hand and what would be needed in the way of theory and data to justify the claims of group differences made by hereditarian scientists on the other. The same applies to some extent to other traditions in the study of human variation. Building a genuinely evolutionarily informed science of human variation gives us the opportunity to re-frame the now stagnant and sterile nature vs. nurture debate on new terms and the present day hereditarian vs. non-hereditarian divide would likely dissolve and be replaced by a new set of more productive questions.

## Acknowledgement

Insert the Acknowledgment text here.

## Funding Statement

This research was supported by an NSF IOS grant 2208944 to KAB; ¡fundername¿ ¡doi¿ (¡award ID¿).

## Competing Interests

None

## Data Availability Statement

All data used are in the tables in the paper and the R code used to produce graphs and model results are available at ****.

## Appendix 1. The Lush-DeFries equation in terms of the coalescent

This is the entire Lush-DeFries equation written in terms of the coalescent prior to any simplification.

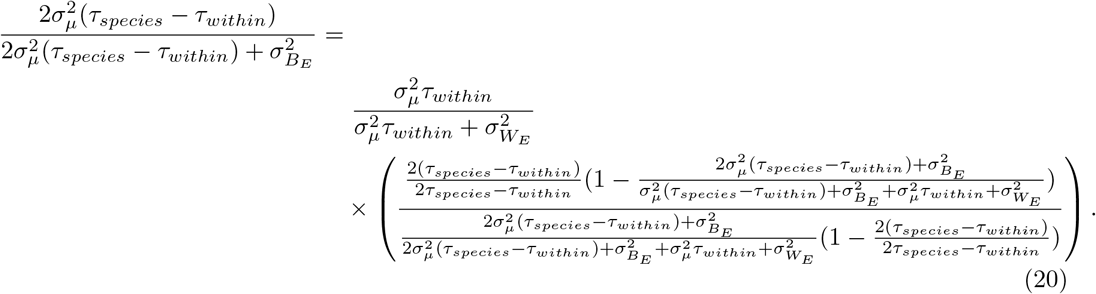

## Appendix 2. Magnitude of differentiation under random genetic drift

Reckoning all evolution with respect to the phenotypic mean of the ancestral population and centering the distribution of evolutionary outcomes such that the ancestral state *E*[*x*_*a*_] = 0, under random genetic drift, *E*[*D*_(*i*,*j*)_] = 0 because *E*[*x*_*i*_− *x*_*j*_] = *E*[*x*_*i*_] −*E*[*x*_*j*_] = 0. However, considerable focus in the hereditarian race science literature is on understanding the absolute number of, say, IQ points separating socially defined racial groups in the United States (Jensen 1998; Warne 2021). If we focus on *E* [|*x*_*i*_ −*x*_*j*_|] for which the sign of the difference between the phenotypic states of the *i*^*th*^ and *j*^*th*^ populations is disregarded, a different picture emerges. Over evolutionary time scales, we can expect the distribution of neutral evolutionary outcomes of a set of lineages evolving independently from a common ancestor at time *τ* = 0 to take the form 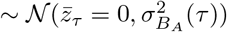. Depending on the application, we can use the different expressions for 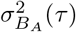 presented above.

The expected absolute difference between two populations, 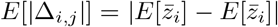 may be obtained using the second L-moment (*λ*_2_) of the (normal) distribution of evolutionary outcomes, which is

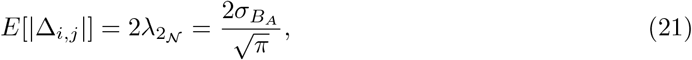

where 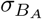 is the among-group additive genetic standard deviation (Bird 2021; Jensen 1998). The absolute difference between pairs of evolutionary outcomes conforms to a slightly modified half-normal distribution. Its variance is given by

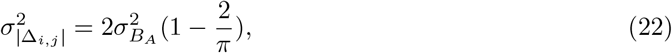

its probability density function by

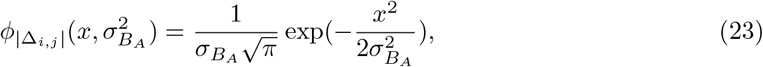

and its cumulative density function is

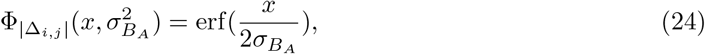

where erf refers to the error function. Substituting the relevant expressions for 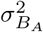 (see section *A neutral evolutionary genetic perspective on* 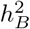) into these expressions allows us to calculate the expectations and variances of evolutionary outcomes and perform basic statistical hypothesis testing on the natural scale of the trait in question.

## Footnotes

1. The preferred nomenclature here depends on the disciplinary context. The psychology literature tends to use “between” as opposed to “among.” Since, in the general case, we would be dealing with more than two groups or families, the more appropriate term is “among” but the literature tends to use the “B” subscript to indicate the among-group heritability (Walsh and Lynch 2018, p.729).

2. The issues surrounding polygenic scores and admixture have already been covered in some depth with respect to the hereditarian hypothesis (Freese et al. 2019; Bird 2021; Bird, Jackson, and Winston, in press, e.g.) but the hereditarian study of the relationship between within- and between- population variation has not been subject to much evolutionary scrutiny for half a century (except see Schraiber and Edge 2023).

3. Warne summarized this strategy as follows: Warne (2021) elaborates on this point: “anyone who postulates that between-group heritability is less than within-group heritability must expect a larger mean environmental difference between groups than the mean difference observed in the actual phenotype. Additionally, the lower the postulated 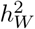 value, the larger environmental differences must be to produce the observed mean phenotype difference… Applied to intergroup intelligence differences, this implies that for between-group differences to be entirely genetic (i.e., for 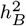 to be zero), the mean between-group environmental differences must be larger than the *d* = 1.00 mean difference observed between African American and White American individuals. Empirical data about the size of these differences can help judge the plausibility of different 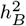 values, including whether 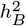 could be equal to zero.” (Warne 2021, p.8)

4. We draw on the following sources for heritabilites: Age at menarche (Towne et al. 2005); age at menopause (Murabito et al. 2005); stature (Roberts, Billewicz, and McGregor 1978); birth weight (Lunde et al. 2007). We use these studies as they estimate heritabilites using non-twin study designs because twin and other sibling based strategies tend to produce inflated estimates of heritability. We found rates of secular change in the following sources: Age at menarche (Jones et al. 2009); age at menopause (Nichols et al. 2006); stature (Khanna and Kapoor 2004); birth weight (Skjaerven, Gjessing, and Bakketeig 2000). These values were selected across genetic and developmental studies such that the secular trends were on the slower end of the spectrum of possibilities and the estimates of *h*2 were collected on groups similar to those from which the secular trend data were drawn.

5. Birth weight has an secular change scaled by 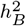 that is an order of magnitude lower than the remainder of the traits, which is not at all evident from the comparison of *d*_*E*_ values. The change in birth weight is not directly comparable to stature, age at menarche, and age at menopause because birth weight is effectively a three-dimensional trait (holding density constant) while age at menarche, age at menopause, and stature are of single dimension and thus have fewer degrees of freedom over which they might vary.

## Notes

### Competing Interest Statement

The authors have declared no competing interest.

## References

1000 Genomes Project Consortium et al. 2015. A global reference for human genetic variation. Nature 526 (7571): 68. 10.1038/nature15393.

Alcala, Nicolas, and Noah A Rosenberg. 2017. Mathematical constraints on FST : Biallelic markers in arbitrarily many populations. Genetics 206, no. 3 (July): 1581–1600. issn: 1943-2631. 10.1534/genetics.116.199141. eprint: https://academic.oup.com/genetics/article-pdf/206/3/1581/42158117/genetics1581.pdf.

Anholt, Robert RH, and Trudy FC Mackay. 2009. Principles of behavioral genetics. Academic Press.

Atchley, William R, and Jun Zhu. 1997. Developmental quantitative genetics, conditional epigenetic variability and growth in mice. Genetics 147 (2): 765–776. 10.1093/genetics/147.2.765.

Berg, Jeremy J, and Graham Coop. 2014. A population genetic signal of polygenic adaptation. PLoS Genetics 10 (8): e1004412. 10.1371/journal.pgen.1004412.

Bird, Kevin A. 2021. No support for the hereditarian hypothesis of the black–white achievement gap using polygenic scores and tests for divergent selection. American Journal of Physical Anthropology 175 (2): 465–476. 10.1002/ajpa.24216.

Bird, Kevin A, John P Jackson, and Andrew S Winston. in press. Confronting scientific racism in psychology: Lessons from evolutionary biology and genetics. American Psychologist.

Bookstein, Fred L. 2016. The inappropriate symmetries of multivariate statistical analysis in geometric morphometrics. Evolutionary Biology 43:277–313.

Carl, Noah. 2018. How stifling debate around race, genes and IQ can do harm. Evolutionary Psychological Science 4 (4): 399–407.

Cheesman, Rosa, Avina Hunjan, Jonathan RI Coleman, Yasmin Ahmadzadeh, Robert Plomin, Tom A McAdams, Thalia C Eley, and Gerome Breen. 2020. Comparison of adopted and nonadopted individuals reveals gene– environment interplay for education in the UK Biobank. Psychological Science 31 (5): 582–591.

Chipuer, Heather M, Michael J Rovine, and Robert Plomin. 1990. LISREL modeling: Genetic and environmental influences on IQ revisited. Intelligence 14 (1): 11–29. 10.1016/0160-2896(90)90011-H.

Cloninger, C Robert, John Rice, and Theodore Reich. 1979. Multifactorial inheritance with cultural transmission and assortative mating. II. a general model of combined polygenic and cultural inheritance. American Journal of Human Genetics 31 (2): 176–198.

Davies, Gail, Albert Tenesa, Antony Payton, Jian Yang, Sarah E Harris, David Liewald, Xiayi Ke, Stephanie Le Hellard, Andrea Christoforou, Michelle Luciano, et al. 2011. Genome-wide association studies establish that human intelligence is highly heritable and polygenic. Molecular Psychiatry 16 (10): 996–1005. 10.1038/mp.2011.85.

DeFries, JC. 1972. Genetics, environment and behavior: implications for educational policy. Chap. Quantitative aspects of genetics and environment in the determination of behavior. Edited by L Ehrman, GS Omenn, and E Caspari. Academic Press. 10.1016/B978-0-12-233450-4.50009-4.

Devlin, Bernie, Michael Daniels, and Kathryn Roeder. 1997. The heritability of IQ. Nature 388 (6641): 468–471. 10.1038/41319.

Ding, Yi, Kangcheng Hou, Ziqi Xu, Aditya Pimplaskar, Ella Petter, Kristin Boulier, Florian Privé, Bjarni J Vilhjálmsson, Loes M Olde Loohuis, and Bogdan Pasaniuc. 2023. Polygenic scoring accuracy varies across the genetic ancestry continuum. Nature, 1–8.

Domingue, Ben. 2014. Evaluating the equal-interval hypothesis with test score scales. Psychometrika 79:1–19.

Downes, Stephen M, and Eric Turkheimer. 2021. An early history of the heritability coefficient applied to humans (1918–1960). Biological Theory, 1–12. 10.1007/s13752-021-00392-9.

Durvasula, Arun, and Alkes Price. 2023. Distinct explanations underlie gene-environment interactions in the UK Biobank. medRxiv, 2023–09.

Edge, Michael D., and Noah A. Rosenberg. 2015. Implications of the apportionment of human genetic diversity for the apportionment of human phenotypic diversity. Genomics and Philosophy of Race, Studies in History and Philosophy of Science Part C: Studies in History and Philosophy of Biological and Biomedical Sciences 52:32–45. issn: 1369-8486. 10.1016/j.shpsc.2014.12.005. https://www.sciencedirect.com/science/article/pii/S1369848615000023.

Falconer, DS, and TFC Mackay. 1996. Introduction to Quantitative Genetics. 3rd. Harlow, Essex, UK: Longmans Green.

Feldman, Marcus W, and Richard C Lewontin. 1975. The heritability hang-up: The role of variance analysis in human genetics is discussed. Science 190 (4220): 1163–1168. https://doi.org/doi/pdf/10.1126.

Feldman, Marcus W, and Richard C Lewontin. 1976. Response: Heritability of IQ. Science 194 (4260): 12–14.

Fogarty, Laurel, and Michael J Wade. 2022. Niche construction in quantitative traits: Heritability and response to selection. Proceedings of the Royal Society B 289 (1976): 20220401.

Frankel, Joseph. 1976. Heritability of IQ. Science 194 (4260): 12–12.

Freese, Jeremy, Ben Domingue, Sam Trejo, Kamil Sicinski, and Pamela Herd. 2019. Problems with a causal interpretation of polygenic score differences between Jewish and non-Jewish respondents in the Wisconsin Longitudinal Study. SocArXiv.

Fuerst, John GR, Emil OW Kirkegaard, and Davide Piffer. 2021. More research needed: There is a robust causal vs. confounding problem for intelligence-associated polygenic scores in context to admixed American populations. Mankind Quarterly 62 (1).

Fuerst, John GR, Vladimir Shibaev, and Emil OW Kirkegaard. 2023. A genetic hypothesis for American race/ethnic differences in mean g: a reply to Warne (2021) with fifteen new empirical tests using the ABCD dataset. Mankind Quarterly 63 (4).

Guo, Jing, Yang Wu, Zhihong Zhu, Zhili Zheng, Maciej Trzaskowski, Jian Zeng, Matthew R Robinson, Peter M Visscher, and Jian Yang. 2018. Global genetic differentiation of complex traits shaped by natural selection in humans. Nature Communications 9 (1): 1–9. 10.1038/s41467-018-04191-y.

Hansen, Thomas F, Christophe Pélabon, and David Houle. 2011. Heritability is not evolvability. Evolutionary Biology 38:258–277.

Hardy, Godfrey H. 1908. Mendelian proportions in a mixed population. Science 28 (706): 49–50. https://doi.org/doi/pdf/10.1126/.

Holland, Dominic, Oleksandr Frei, Rahul Desikan, Chun-Chieh Fan, Alexey A Shadrin, Olav B Smeland, Vijay S Sundar, Paul Thompson, Ole A Andreassen, and Anders M Dale. 2020. Beyond SNP heritability: polygenicity and discoverability of phenotypes estimated with a univariate Gaussian mixture model. PLoS Genetics 16 (5): e1008612. 10.1371/journal.pgen.1008612.

Howe, Laurence J, Michel G Nivard, Tim T Morris, Ailin F Hansen, Humaira Rasheed, Yoonsu Cho, Geetha Chittoor, Rafael Ahlskog, Penelope A Lind, Teemu Palviainen, et al. 2022. Within-sibship genome-wide association analyses decrease bias in estimates of direct genetic effects. Nature genetics 54 (5): 581–592.

Hu, Sile, Lino AF Ferreira, Sinan Shi, Garrett Hellenthal, Jonathan Marchini, Daniel J Lawson, and Simon R Myers. 2023. Leveraging fine-scale population structure reveals conservation in genetic effect sizes between human populations across a range of human phenotypes. bioRxiv, 2023–08.

Jackson, John P, and Andrew S Winston. 2021. The mythical taboo on race and intelligence. Review of General Psychology 25 (1): 3–26. 10.1177/2F1089268020953622.

Jensen, Arthur R. 1969. How much can we boost IQ and educational achievement. Harvard Educational Review 39 (1): 1–123.

Jensen, Arthur R.. 1974. Race and mental ability. In Symposium of the Institute of Biology on “Racial Variation in Man”. Royal Geographical Society.

Jensen, Arthur R.. 1975. The meaning of heritability in the behavioral sciences. Educational Psychologist 11 (3): 171–183. 10.1080/00461527509529142.

Jensen, Arthur R. 1976. Heritability of IQ. Science 194 (4260): 6–8.

Jensen, Arthur R.. 1998. The g factor: The science of mental ability. Westport, Connecticut: Praeger.

Jones, Laura L., Paula L. Griffiths, Shane A. Norris, John M. Pettifor, and Nöel Cameron. 2009. Age at menarche and the evidence for a positive secular trend in urban South Africa. American Journal of Human Biology 21 (1): 130–132. 10.1002/ajhb.20836. eprint: https://onlinelibrary.wiley.com/doi/pdf/10.1002/ajhb.20836.https://onlinelibrary.wiley.com/doi/abs/10.1002/ajhb.20836.

Kanazawa, Satoshi. 2008. Temperature and evolutionary novelty as forces behind the evolution of general intelligence. Intelligence 36 (2): 99–108. 10.1016/j.intell.2007.04.001.

Khanna, Geeta, and Satwanti Kapoor. 2004. Secular trend in stature and age at menarche among Punjabi Aroras residing in New Delhi, India. Collegium Antropologicum 28 (2): 571–575.

Kim, Jaehee, Michael D. Edge, Amy Goldberg, and Noah A. Rosenberg. 2021. Skin deep: The decoupling of genetic admixture levels from phenotypes that differed between source populations. American Journal of Physical Anthropology 175 (2): 406–421. 10.1002/ajpa.24261. eprint: https://onlinelibrary.wiley.com/doi/pdf/10.1002/ajpa.24261.https://onlinelibrary.wiley.com/doi/abs/10.1002/ajpa.24261.

Lande, Russell. 1976. Natural selection and random genetic drift in phenotypic evolution. Evolution, 314–334. https://doi.org/doi.org/2407703.

Lasker, Jordan, Bryan J. Pesta, John G. R. Fuerst, and Emil O. W. Kirkegaard. 2019. Global ancestry and cognitive ability. Psych 1 (1): 431–459. issn: 2624-8611. 10.3390/psych1010034.

Lewontin, Richard C. 1970. Race and intelligence. Bulletin of the Atomic Scientists 26 (3): 2–8. 10.1080/00963402.1970.11457774.

Loehlin, John C. 1978. Heredity-environment analyses of Jencks’s IQ correlations. Behavior Genetics 8 (5): 415–436. 10.1007/BF01067938.

Lunde, Astrid, Kari Klungsøyr Melve, Håkon K Gjessing, Rolv Skjærven, and Lorentz M Irgens. 2007. Genetic and environmental influences on birth weight, birth length, head circumference, and gestational age by use of population-based parent-offspring data. American Journal of Epidemiology 165 (7): 734–741.

Lush, Jay L. 1947a. Family merit and individual merit as bases for selection. Part I. The American Naturalist 81 (799): 241–261. 10.1086/281520.

Lush, Jay L.. 1947b. Family merit and individual merit as bases for selection. Part II. The American Naturalist 81 (800): 362–379. 10.1086/281520.

Lynch, Michael, and Bruce Walsh. 1998. Genetics and Analysis of Quantitative Traits. Vol. 1. Sinauer Sunderland, MA.

Lynn. 2019. Reflections on sixty-eight years of research on race and intelligence. Psych 1 (1): 123–131.

Lynn, Richard. 1991. The evolution of racial differences in intelligence. Mankind Quarterly 32 (1): 109.

Lynn, Richard. 2008. The global bell curve: race, IQ, and inequality worldwide. Washington Summit Publishers.

Miller, Geoffrey F, and Lars Penke. 2007. The evolution of human intelligence and the coefficient of additive genetic variance in human brain size. Intelligence 35 (2): 97–114.

Morton, Newton E. 1976. Heritability of IQ. Science 194 (4260): 9–10.

Mostafavi, Hakhamanesh, Arbel Harpak, Ipsita Agarwal, Dalton Conley, Jonathan K Pritchard, and Molly Przeworski. 2020. Variable prediction accuracy of polygenic scores within an ancestry group. eLife 9:e48376.

Muir, William M. 1996. Group selection for adaptation to multiple-hen cages: Selection program and direct responses. Poultry Science 75 (4): 447–458. 10.3382/ps.0750447.

Murabito, Joanne M, Qiong Yang, Caroline Fox, Peter WF Wilson, and L Adrienne Cupples. 2005. Heritability of age at natural menopause in the Framingham Heart Study. The Journal of Clinical Endocrinology & Metabolism 90 (6): 3427–3430.

Nichols, Hazel B, Amy Trentham-Dietz, John M Hampton, Linda Titus-Ernstoff, Kathleen M Egan, Walter C Willett, and Polly A Newcomb. 2006. From menarche to menopause: trends among US women born from 1912 to 1969. American Journal of Epidemiology 164 (10): 1003–1011.

Panofsky, Aaron, Kushan Dasgupta, and Nicole Iturriaga. 2021. How white nationalists mobilize genetics: From genetic ancestry and human biodiversity to counterscience and metapolitics. American Journal of Physical Anthropology 175 (2): 387–398.

Plomin, Robert, and John C DeFries. 1976. Heritability of IQ. Science 194 (4260): 10–12.

Polderman, Tinca JC, Beben Benyamin, Christiaan A De Leeuw, Patrick F Sullivan, Arjen Van Bochoven, Peter M Visscher, and Danielle Posthuma. 2015. Meta-analysis of the heritability of human traits based on fifty years of twin studies. Nature Genetics 47 (7): 702. 10.1038/ng.3285.

Postma, Erik. 2014. Four decades of estimating heritabilities in wild vertebrate populations: Improved methods, more data, better estimates. In Quantitative genetics in the wild, edited by Anne Charmantier, Dany Garant, and Loeske EB Kruuk, 33. OUP Oxford.

Racimo, Fernando, Jeremy J Berg, and Joseph K Pickrell. 2018. Detecting polygenic adaptation in admixture graphs. Genetics 208 (4): 1565–1584. 10.1534/genetics.117.300489.

Rao, DC, NE Morton, JM Lalouel, and R Lew. 1982. Path analysis under generalized assortative mating: II. American IQ. Genetics Research 39 (2): 187–198. doi:10.1017/S0016672300020875.

Refoyo-Martínez, Alba, Siyang Liu, Anja Moltke Jørgensen, Xin Jin, Anders Albrechtsen, Alicia R Martin, and Fernando Racimo. 2021. How robust are cross-population signatures of polygenic adaptation in humans? Peer Community Journal 1.

Roberts, D. F., W. Z. Billewicz, and I. A. McGregor. 1978. Heritability of stature in a West African population. Annals of Human Genetics 42 (1): 15–24. 10.1111/j.1469-1809.1978.tb00928.x. eprint: https://onlinelibrary.wiley.com/doi/pdf/10.1111/j.1469-1809.1978.tb00928.x.https://onlinelibrary.wiley.com/doi/abs/10.1111/j.1469-1809.1978.tb00928.x.

Rushton, J Philippe. 1985. Differential K theory: the sociobiology of individual and group differences. Personality and Individual Differences 6 (4): 441–452. 10.1016/0191-8869(85)90137-0.

Rushton, J Philippe. 1995. Race, evolution, and behavior: A life history perspective. Transaction.

Rushton, J Philippe, and Arthur R Jensen. 2005. Thirty years of research on race differences in cognitive ability. Psychology, Public Policy, and Law 11 (2): 235. 10.1037/1076-8971.11.2.235.

Schraiber, Joshua G., and Michael D. Edge. 2023. Heritability within groups is uninformative about differences among groups: Cases from behavioral, evolutionary, and statistical genetics. bioRxiv, 10.1101/2023.11.06.565864. eprint: https://www.biorxiv.org/content/early/2023/11/06/2023.11.06.565864.full.pdf.https://www.biorxiv.org/content/early/2023/11/06/2023.11.06.565864.

Sear, Rebecca. 2022. ‘national IQ’datasets do not provide accurate, unbiased or comparable measures of cognitive ability worldwide.

Sesardic, Neven. 2000. Philosophy of science that ignores science: Race, IQ and heritability. Philosophy of Science 67 (4): 580–602.

Sesardic, Neven. 2005. Making sense of heritability. Cambridge University Press.

Skjaerven, Rolv, Håkon K Gjessing, and LEIV S Bakketeig. 2000. Birthweight by gestational age in Norway. Acta Obstetricia et Gynecologica Scandinavica 79 (6): 440–449.

Slatkin, Montgomery. 1991. Inbreeding coefficients and coalescence times. Genetics Research 58 (2): 167–175.

Stock, Jay. 2023. The extended evolutionary synthesis and distributed adaptation in the genus Homo: Phenotypic plasticity and behavioral adaptability. PaleoAnthropology 2023 (2): 205–233.

Towne, Bradford, Stefan A Czerwinski, Ellen W Demerath, John Blangero, Alex F Roche, and Roger M Siervogel. 2005. Heritability of age at menarche in girls from the Fels Longitudinal Study. American Journal of Physical Anthropology 128 (1): 210–219.

Uchiyama, Ryutaro, Rachel Spicer, and Michael Muthukrishna. 2022. Cultural evolution of genetic heritability. Behavioral and Brain Sciences 45:e152.

Voje, Kjetil Lysne, James G Saulsbury, Jostein Starrfelt, Daniel Varajão Latorre, Alexis Rojas, Vilde Bruhn Kinneberg, Lee Hsiang Liow, Connor J Wilson, Erin E Saupe, and Mark Grabowski. 2023. Measurement theory and paleobiology. Trends in Ecology & Evolution.

Walsh, Bruce, and Michael Lynch. 2018. Evolution and Selection of Quantitative Traits. Oxford University Press.

Warne, Russell T. 2020. In the know: Debunking 35 myths about human intelligence. Cambridge University Press.

Warne, Russell T.. 2021. Between-group mean differences in intelligence in the United States are > 0% genetically caused: five converging lines of evidence. The American Journal of Psychology 134 (4): 480–501. 10.5406/amerjpsyc.134.4.0479.

Warne, Russell T, Jared Z Burton, Aisa Gibbons, and Daniel A Melendez. 2019. Stephen Jay Gould’s analysis of the Army Beta test in The Mismeasure of Man: Distortions and misconceptions regarding a pioneering mental test. Journal of Intelligence 7 (1): 6.

Whitlock, Michael C. 1999. Neutral additive genetic variance in a metapopulation. Genetics Research 74 (3): 215–221.

Whitlock, Michael C.. 2008. Evolutionary inference from QST . Molecular Ecology 17 (8): 1885–1896. 10.1111/j.1365-294X.2008.03712.x.

Whitlock, Michael C, and Frederic Guillaume. 2009. Testing for spatially divergent selection: Comparing QST to FST . Genetics 183 (3): 1055–1063. 10.1534/genetics.108.099812.

Winegard, Bo, Ben Winegard, and Jonathan Anomaly. 2020. Dodging Darwin: Race, evolution, and the hereditarian hypothesis. Personality and Individual Differences 160:109915. 10.1016/j.paid.2020.109915.

Yair, Sivan, and Graham Coop. 2022. Population differentiation of polygenic score predictions under stabilizing selection. Philosophical Transactions of the Royal Society B 377 (1852): 20200416.

Young, Alexander I, Michael L Frigge, Daniel F Gudbjartsson, Gudmar Thorleifsson, Gyda Bjornsdottir, Patrick Sulem, Gisli Masson, Unnur Thorsteinsdottir, Kari Stefansson, and Augustine Kong. 2018. Relatedness disequilibrium regression estimates heritability without environmental bias. Nature Genetics 50 (9): 1304–1310. 10.1038/s41588-018-0178-9.

Zeng, Jian, Ronald De Vlaming, Yang Wu, Matthew R Robinson, Luke R Lloyd-Jones, Loic Yengo, Chloe X Yap, Angli Xue, Julia Sidorenko, Allan F McRae, et al. 2018. Signatures of negative selection in the genetic architecture of human complex traits. Nature Genetics 50 (5): 746–753.

